# A role for HDAC3 in regulating histone lactylation and maintaining oocyte chromatin architecture and fertility

**DOI:** 10.64898/2026.01.26.701805

**Authors:** Inês Simões-Gomes, António Jacinto, Ana Pimenta-Marques

## Abstract

The establishment of specialized chromatin architecture is critical for oocyte development, meiotic fidelity, and embryogenesis. Histone lysine lactylation (Kla) has recently emerged as a widespread chromatin modification, yet its developmental dynamics and function in the germ line remain unclear. Here, we define the temporal profile of Kla during *Drosophila* oogenesis and identified enzymes that regulate its levels. Kla is absent in early germarial stages, becomes highly enriched during prophase I as meiotic chromosomes condense into the karyosome, and decreases when the karyosome decompacts during transcriptional reactivation. Germline depletion of HDAC3 and Nej/p300 reduces Kla in stage-10 oocytes, revealing conserved chromatin-modifying enzymes that maintain lactylation. Among these, HDAC3 is essential for sustaining karyosome architecture, assembling a normal meiotic spindle, and supporting fertility. These findings identify histone lactylation as a dynamic chromatin feature of oogenesis and suggest that Kla contributes to the establishment of meiotic chromatin states required for oocyte competence.

**SUMMARY STATEMENT:** Histone lactylation is developmentally regulated during *Drosophila* oogenesis. HDAC3 maintains histone lactylation and is essential for meiotic chromatin architecture, meiotic progression, and female fertility.

## INTRODUCTION

Proper gamete formation is essential for the continuity of life and for ensuring successful embryonic development. In animal oocytes, chromatin organization plays a central role in meiotic progression, transcriptional control, and the acquisition of developmental competence^1,2^. Transcriptional silencing of the oocyte nucleus for extended periods is a conserved feature across metazoans, occurring at species-specific stages. During these stages, the oocyte nucleus chromatin is maintained in a highly compacted state. Disruption of this state with premature chromatin decompaction compromises meiotic fidelity and reduces fertility^1,2^, underscoring the tight coupling between chromatin structure and oocyte function.

An important regulator of chromatin architecture and gene expression is the epigenetic modification of histones. Histone lysine post-translational modifications (PTMs), such as acetylation and methylation, play critical roles in modulating chromatin compaction and transcriptional states^3,4^. Multiple PTMs, including acetylation, methylation and phosphorylation have been described in developing oocytes, implying that epigenetic modifications play important roles during oocyte maturation^5–8^. In oocytes, these epigenetic modifications coordinate nuclear organization, ensure accurate meiotic progression, and are key determinants of developmental competence^2,6,7,9^. Work in *Drosophila* showed that perturbing the oocyte epigenome alters chromatin compaction, transcriptional programs and leads to meiotic defects^7^. Despite these advances, the mechanisms that establish, interpret and dynamically remodel the oocyte epigenome are still not fully elucidated, reflecting the high complexity and plasticity of these regulatory pathways.

Recent work identified histone lysine lactylation (Kla) as a novel PTM derived from lactate, a product of glycolysis^10^, and which has been associated to transcriptional activation^10,11^. This modification has been detected across a broad evolutionary range, from plants to mammals^10,12–14^, and has been implicated in diverse biological processes such as tumor progression^15^, regulation of inflammation^16^ and neuronal function^17^. Very recently, histone lactylation was detected in the nuclei of oocytes and sperm from both mouse^14,18,19^ and *Drosophila*^*20*^, suggesting that this modification may play conserved roles in gametogenesis across species. In *Drosophila*, histone lactylation was reported predominantly at stages of oogenesis characterized by transcriptionally silencing^*20*^, when the chromatin is highly compacted. These observations suggest that lactylation may contribute to additional unknown functions beyond its established role in transcriptional activation. However, it remains unclear how histone lactylation correlates with different stages of oocyte maturation and whether it plays a physiological relevant role throughout oogenesis.

To address these questions, we systematically characterized Kla throughout *Drosophila* oogenesis. Quantitative analyses revealed that Kla levels in the oocyte are dynamically regulated during oogenesis. Histone lactylation is absent or markedly reduced in the germarium, peaking at stages when the oocyte assembles the highly compacted karyosome chromatin, and decreases again as the karyosome decompacts in later stages. Using a RNAi-based approach, we identified chromatin modifying enzymes, including HDAC3, as regulators of Kla and chromosome architecture. Depletion of HDAC3 in the oocyte led to reduced lactylation, premature chromatin decompaction, meiotic defects, and decreased fertility. Together, these observations reveal that histone lactylation is a dynamic chromatin modification in the oocyte and identify HDAC3 as a key regulator of both Kla levels and oocyte chromosome architecture.

## RESULTS AND DISCUSSION

### Histone lysine lactylation is dynamically regulated during oogenesis and decreases upon natural chromatin architecture remodeling

In *Drosophila*, oocyte transcription becomes globally silenced at stage 5 and remains inactive through stage 9 of oogenesis^7,21^ (Fig.1A) This transcriptional quiescence coincides with the reorganization of the oocyte chromatin into a highly compacted cluster of meiotic chromosomes known as the karyosome^22,23^, a hallmark of prophase I arrest. Uridine-incorporation assays have shown that the oocyte nucleus is transcriptionally active from stages 9-10 of oogenesis, which occurs concomitantly with a partial decompaction of the karyosome DNA^7,21^. Given these well-defined nuclear transitions, we took advantage of the distinct morphological and transcriptional stages of *Drosophila* oogenesis to investigate how Kla levels change during oocyte development.

**Figure 1:**
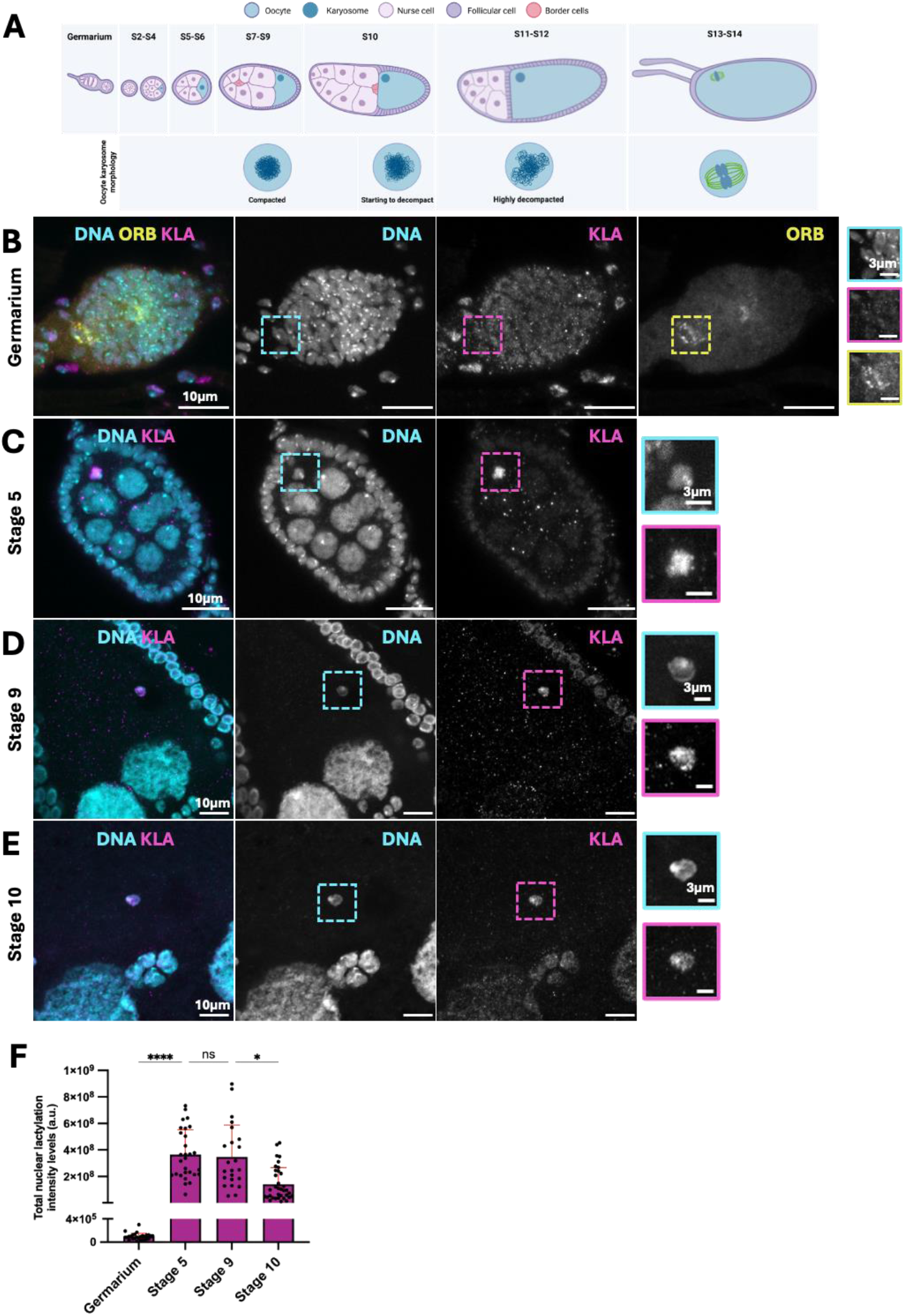
Histone lactylation at the *Drosophila* oocyte is developmentally regulated. **(A)** Schematic of Drosophila oogenesis. In *Drosophila*, each ovary is composed on average by 16 ovarioles, which are strings of progressively developing egg chambers. Oogenesis begins at the tip of a structure called the germarium, with the asymmetric division of a germ-line stem cell to give rise to a new stem cell and a cytoblast, which undergoes four successive mitotic cycles with incomplete cytokinesis, forming a large cyst composed of 16 interconnected cells. At late stage 2a of the germarium, one of the 16 cells becomes the oocyte, while the other 15 cells take on the role of nurse cells, which generate proteins and mRNAs that are provided to the developing oocyte. The specified oocyte and its associated nurse cells are then encapsulated by somatic cells to form an egg chamber that buds off from the germarium. Created in https://BioRender.com. **(B-D)** Representative immunofluorescence images showing distinctive patterns of histone lactylation in *Drosophila* oocytes. Egg chambers were immunostained for histone lysin lactylation (Kla, magenta), Orb (yellow) and stained for DNA (cyan). Merged channels are shown alongside individual channels for each marker. Insets show higher magnification views of oocyte nuclei. Scale bars: 10µm and 3µm (insets). **(E)** Temporal analysis of the dynamic levels of Kla throughout oogenesis. Relative levels are expressed in fluorescence arbitrary units (a.u.). Each dot represents one oocyte nucleus; mean ± SEM are shown. Statistical analysis was performed using one-way ANOVA followed by Kruskal-Wallis’ multiple comparisons test. ****p < 0.0001, *p < 0.01, ns = not significant. N = 3, minimum of 10 oocytes per experiment.

While prior work reported the presence of Kla and its enrichment in the oocyte nucleus at mid oogenesis^20^, its temporal dynamics relative to chromatin architecture has not been quantified. To address this gap, we performed a quantitative assessment of histone lactylation to determine how Kla levels fluctuate as oocytes progress through these key nuclear transitions. Using a pan-lactylation antibody, we quantified Kla in the oocyte nucleus at four representative stages, which included the germarium, the stage at which the oocyte is determined within cysts of 16 cells, and stages 5, 9 and 10 of oogenesis (Fig.1 A)^24^. To identify oocytes in the germarium, we used an antibody against Orb, a cytoplasmic RNA-binding protein that marks the developing oocyte shortly after specification in the germarium^24^. No detectable enrichment of Kla was observed in Orb-positive germarial oocytes (Fig.1B), indicating that Kla levels were very low or absent at early stages of oocyte specification. Kla signal first appeared at stage 3 (Supp. Fig.1A) and became strongly enriched in the oocyte nucleus from stages 5 to 9, after which it appeared to decrease at stages 10 (Fig.1C-F). Quantification of Kla levels confirmed that lactylation increases sharply from the germarium to stages 4 of oogenesis. Kla levels at stages 4 and 9 were comparable, suggesting that histone lactylation remains stable throughout the mid stages of oogenesis, and then decreases at stages 10 (Fig. 1B-F), coinciding with the onset of chromatin decompaction and transcriptional reactivation^7,21^.

Our analysis shows that histone lactylation is developmentally regulated during oogenesis, with high levels maintained during the transcriptionally inactive, compact chromatin stages, and a clear decline at stage 10, coinciding with chromatin decompaction. Kla has been proposed to act as a transcription-activating modification, particularly in contexts where glycolytic metabolism is high, such as in cancer^10,25,26^. However, our findings, together with recent observations, challenge this simplified model, as Kla levels peak at stages when meiotic chromosomes are tightly compacted into the karyosome, a chromatin configuration associated with transcriptionally silencing rather than activation. Moreover, we also detected strong lactylation in embryonic nuclei, including during early nuclear divisions when transcription is silenced (Supp. Fig. 3). Syncytial embryo nuclei also display specialized chromatin architectures that support the extremely fast mitotic divisions, characteristic of early embryogenesis^27,28^. Consistent with this, mouse oocytes and early embryos also show dynamic lactylation patterns, with high nuclear Kla at the prophase I germinal vesicle (GV) stage, when chromatin is highly compacted, and a progressive increase in Kla levels from the 2-cell stage to the blastocyst^14,29^.

Together, these observations indicate that dynamic regulation of histone lactylation is a conserved feature of oocyte maturation and early embryogenesis. Rather than functioning solely as a transcriptional activation mark, Kla may promote the establishment or maintenance of specialized chromatin configurations in a stage-specific manner, which may in turn influence transcription. During oogenesis, Kla may promote or reinforce the highly compact karyosome structure characteristic of transcriptionally silent prophase I oocytes. Notably, Kla is absent from the germarium, where transcriptional regulation is essential for germline stem-cell maintenance and oocyte specification^2,30^, suggesting that histone lactylation is unlikely to play a major role in the earliest steps of oocyte determination. Our observations support a model in which histone lactylation contributes to the establishment or maintenance of distinct chromatin architectures in a developmental stage–specific manner, rather than acting exclusively through direct modulation of transcription.

### Multiple conserved chromatin-modifying enzymes are required to maintain histone lysine lactylation in the oocyte

If histone lactylation contributes to the establishment or maintenance of a compacted chromatin state associated with transcriptional silencing during oogenesis, its developmental regulation is likely to depend on enzymes that add and remove lactyl groups in a stage-specific manner. In mammals, several conserved histone acetyltransferases (HATs) and histone deacetylases (HDACs) have been shown to promote histone and non-histone lactylation^25,31,32^. Guided by these observations, we tested whether the *Drosophila* orthologs of these conserved enzymes can function as regulators of histone lactylation in the oocyte. We focused on Nejire (Nej/p300), the *Drosophila* orthologue of p300, and the best characterized histone lactylation writer in mammals^10,18,33,34^ and on HDAC3, a class I deacetylase, implicated in histone delactylation^25,32,35,23^. We performed germline specific *in vivo* RNA interference (RNAi) to individually deplete these candidates and examined whether their loss altered Kla levels in the oocyte karyosome. We specifically focused on stage-10 oocytes, at which lactylation levels naturally decrease as the karyosome begins to decompact (Fig.1F). Depletion of either HDAC3 or Nej resulted in a significant reduction in Kla levels in stage 10 oocytes compared with controls (Fig. 2B-E). For HDAC3, two independent RNAi lines were tested, both of which led to a comparable reduction in karyosome lactylation (Fig. 2C-E). These results show that Nej/p300 promotes histone lactylation in the female germline, consistent with reports that overexpression of p300 in mouse oocytes increases both H4K12 lactylation and acetylation^18^. Notably, loss of HDAC3 also reduced, rather than increased, Kla levels acting in the opposite direction to what would be expected for HDACs functioning solely as erasers of Kla^25,35^. However, consistent with our observations, very recent work demonstrated that class I HDACs (HDAC1-3) can directly catalyze lysine lactylation in mammalian cells at physiological lactate concentrations^32^. Together, these results identify Nej/p300 and HDAC3 as regulators of histone lactylation in the oocyte and suggest that both enzymes contribute to the developmental control of Kla levels during oogenesis.

**Figure 2:**
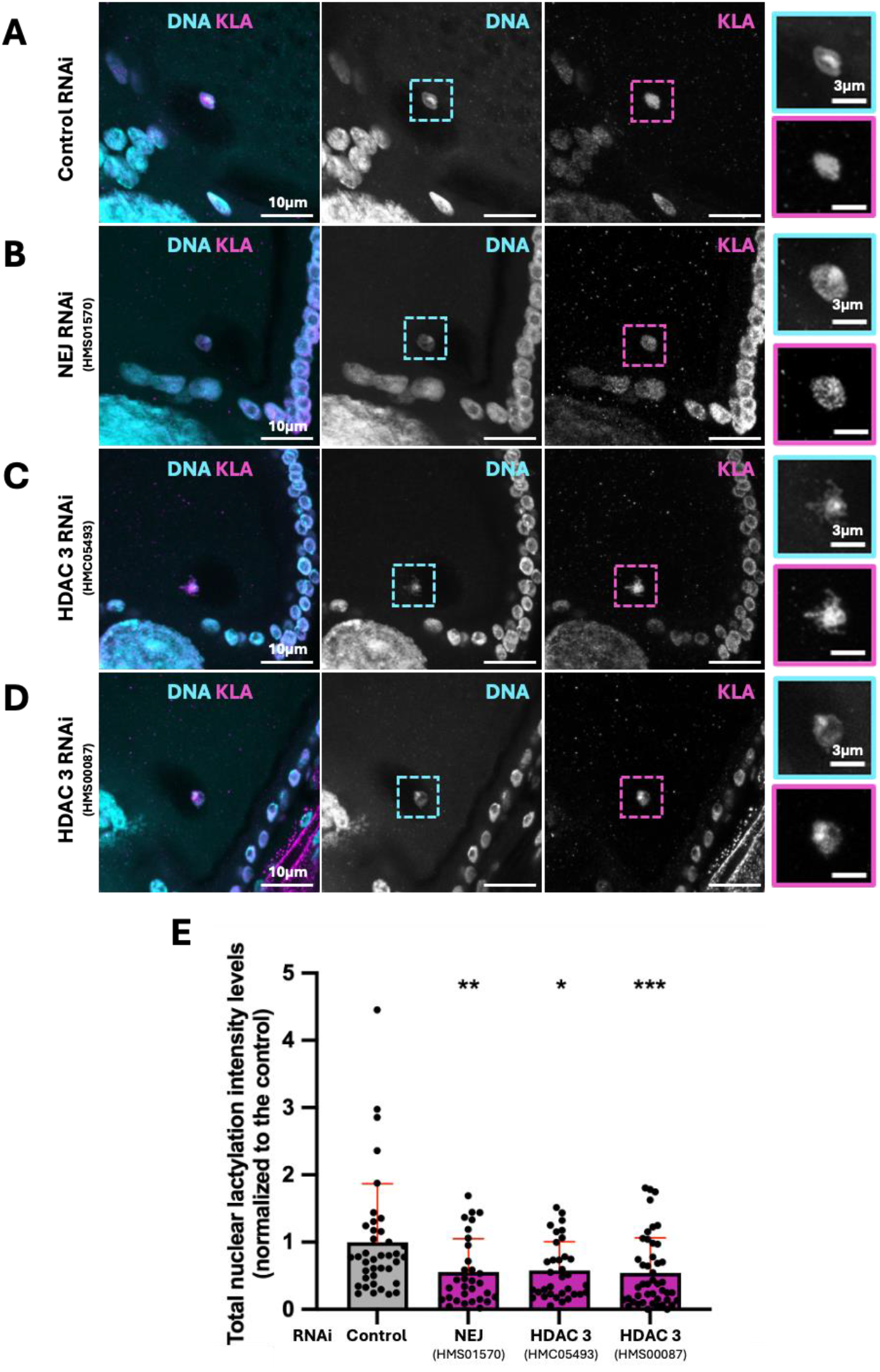
Depletion of Nej and HDAC3 leads to the reduction of histone lysine lactylation in stage-10 oocytes. **(A–D)** Representative immunofluorescence images of stage-10 oocytes from control (*mCherry* RNAi) and RNAi-mediated knockdown of the chromatin modifying proteins Nej and HDAC3, driven by Nanos-Gal4. Kla is shown in magenta and DNA in cyan. Merged images are shown alongside individual channels and higher-magnification views of oocyte nuclei. Scale bars: 10 µm; 3 µm (insets). **(E)** Quantification of Kla levels in stage 10 oocyte nuclei for each condition. Each dot represents one oocyte nucleus; bars indicate mean ± SEM. Statistical analysis was performed using one-way ANOVA followed by Kruskal–Wallis multiple-comparisons testing, with each RNAi condition compared to the control. ***p < 0.001, **p < 0.01, *p < 0.05; ns, not significant. *N* = 3 independent biological replicates, with a minimum of 10 oocytes analyzed per experiment.

### HDAC3 is required to maintain proper karyosome organization

Previous work has shown that chromatin-modifying enzymes are required for maintaining proper meiotic chromatin architecture in the *Drosophila* oocyte^7,36,37^. Because stage-10 oocytes normally transition from a tightly compacted karyosome to a slightly more relaxed chromatin architecture (Fig. 3), we asked whether the enzymes that regulate histone lactylation might influence this remodeling step. Given that depletion of Nej and HDAC3 reduced histone lactylation levels in stage-10 oocytes, we examined whether their loss affected the extent of karyosome remodeling at this stage. We first performed a blind qualitative assessment of karyosome morphology by classifying karyosomes as compact, partially decompacted, or markedly decompacted (Fig. 3A). In control oocytes, the karyosome was predominantly compact, maintaining a smooth and rounded morphology, with 84.2% of oocytes falling into this category (Fig. 3A,C). Nej-depleted oocytes showed karyosome morphologies largely comparable to controls, although a non-significant trend toward decompaction was observed in a subset of oocytes (Fig. 3B,C). Notably, HDAC3-depleted oocytes showed a marked reduction in karyosome compaction. For the HDAC3^*HMC05493*^ RNAi line, only 2.8% of oocytes showed a compact karyosome, while 41.7% were partially decompacted (7,9% in controls) and 55.5% markedly decompacted (7,9% in controls) (Fig. 3B,C). The HDAC3^*HMS00087*^ RNAi line showed a similar, though less pronounced, phenotype, with 33,3% of oocytes displaying compacted chromatin masses (Fig. 3B,C). Together, these observations indicate that depletion of HDAC3 compromises karyosome morphology during stage 10 of oogenesis, whereas depletion of Nej produces minimal or no detectable effect with the RNAi line tested.

**Figure 3:**
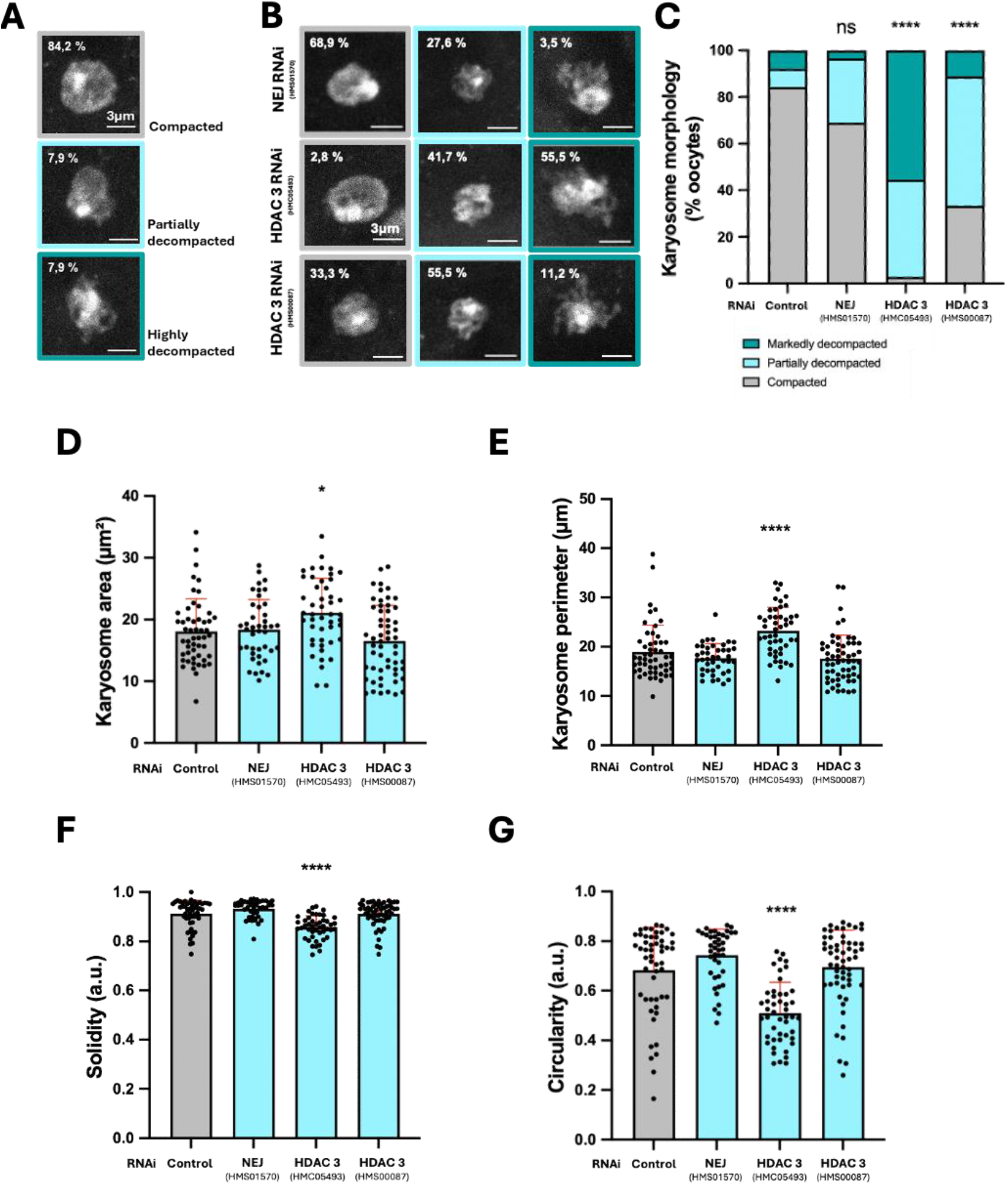
HDAC3 depletion disrupts karyosome organization in stage-10 oocytes. **(A)** Representative images of DAPI-stained nuclei illustrating the three karyosome morphology categories (compact, partially decompacted and highly decompacted) scored in control oocytes (*UAS-mCherry RNAi*), with the corresponding percentages indicated. Scale bar: 3 µm. **(B)** Representative images of DAPI-stained oocyte nuclei following RNAi-mediated knockdown of Nej and HDAC3, with percentages scored for each category. Scale bar: 3 µm. Scale bar: 3 µm. **(C)** Quantification of the percentage of oocytes in each karyosome morphology category. Statistical significance was assessed using Chi-square test comparing the distribution of karyosome categories in each RNAi condition to control. **(D–G)** Quantification of nuclear area (µm^2^), perimeter (µm), circularity (4π × area/perimeter^2^, range 0–1, with 1 = perfect circle), and solidity (area/convex area, range 0–1, with 1 = compact nucleus) across conditions. Each dot represents one oocyte nucleus; mean ± SEM are shown. Statistical analysis was performed using one-way ANOVA followed by Kruskal-Wallis’ multiple comparisons test comparing each RNAi condition to control. **** p < 0.0001, * p < 0.05, ns = not significant. N = 3 independent biological replicates, with a minimum of 10 oocytes analyzed per experiment.

To obtain an objective quantitative assessment of karyosome architecture, we measured several geometric parameters of karyosome structure, including projected area, perimeter, circularity, and solidity (Fig. 2D-G). Circularity quantifies how close an object is to a perfect circle (value of 1), reporting on the overall chromatin mass roundness, whereas solidity measures the ratio of karyosome area to its convex hull, reflecting boundary irregularities. Highly compact, smooth karyosomes are expected to exhibit solidity values closer to 1, whereas irregular or partially fragmented structures are expected to display lower values. By analyzing these parameters in the control oocytes, we found that oocytes classified as “partially” or “markedly” compacted show an increase in the area and perimeter of the karyosome, while the circularity and solidity decreased as compared to the “compacted” oocytes (Supp. Fig. 2A-D). Increases in karyosome perimeter were previously shown to correspond to the expansion of the chromatin mass^7^. Therefore, together, these metrics provide complementary information on karyosome architecture. In controls, most oocytes showed a compacted karyosome with a mean area of 18,1mm^2^ ± 0,7312 SEM and a mean perimeter of 19,0mm ± 0,7599 SEM (Fig. 3D,E). Among all RNAi conditions tested, only oocytes depleted of HDAC3 (using the HDAC3^*HMC05493*^ line) showed statistically significant increases in both parameters, with a mean area of 21,1mm^2^ ± 0,8125 SEM, and a mean perimeter of 23,2 mm ± 0,7015 SEM (Fig. 3D,E). Control karyosomes also exhibited high solidity (0,91± 0,0075 SEM), consistent with their smooth, compact structure, and a circularity of 0.68 ± 0,0239 SEM. Again, only HDAC3^*HMC05493*^ depleted oocytes showed significant reductions, with solidity and circularity values of 0.85 ± 0,0068 SEM and 0.51 ± 0,0180 SEM, respectively (Fig. 3F,G). Neither the HDAC3^*HMSO087*^ RNAi line nor Nej RNAi produced significant changes in these quantitative metrics relative to controls.

Altogether, these results show that loss of HDAC3 disrupts normal karyosome compaction in stage 10 oocytes, indicating that this chromatin modifying enzyme plays an important role in maintaining chromatin architecture at this developmental stage. A conserved feature of metazoan oocytes is that chromatin organization during prophase I is essential for coordinating transcriptional quiescence, ensuring proper meiotic progression and establishing oocyte developmental competence^2,6,7,9.^ Consistent with this, chromatin modifying enzymes such as the kinase Nhk-1^37,38^, and the demethylase kdm5/Lid^7,36^ are required for karyosome assembly and maintenance, as their loss leads to karyosome fragmentation or decompaction. We did not observe any obvious defect in karyosome assembly at earlier stages or karyosome fragmentation, suggesting that HDAC3 is not required for initial karyosome formation. Instead, these results suggest that HDAC3 is required to maintain karyosome integrity during stage 10, when partial decompaction normally occurs alongside transient transcription^7,21^. Nej depletion did not significantly affect karyosome morphology, indicating that Nej is unlikely to be a major regulator of meiotic chromatin remodeling at this stage, although we cannot exclude incomplete RNAi-mediated depletion.

### HDAC3 is required for normal meiosis and female fertility

Conserved chromatin regulators, such as the histone demethylase Kdm5/lid and the histone kinase Nhk-1 are required for karyosome integrity, and their loss leads to meiotic defects and reduced fertility^7,37^. Because HDAC3 depletion resulted in a pronounced karyosome decompaction phenotype, we next asked whether HDAC3 is required for proper meiotic progression. To address this, we examined metaphase I in stage-14 oocytes using Jupiter-GFP (Jup-GFP), a microtubule-binding protein that marks spindle microtubules^39,40^. In control oocytes, the metaphase I spindle typically adopts a stable bipolar, fusiform structure with tightly focused poles. The chromosomes form a compact mass at the equator, where they remain arrested until completion of meiosis after egg activation (Fig. 4C’). In contrast, HDAC3-depleted oocytes frequently displayed spindles with abnormal morphologies. These defects included unfocused poles (Fig. 4C’’) and frayed microtubules (Fig. 4C’’’). These observations suggest that HDAC3 is required for proper meiotic spindle organization and/or the establishment of stable kinetochore–microtubule attachments.

**Figure 4.**
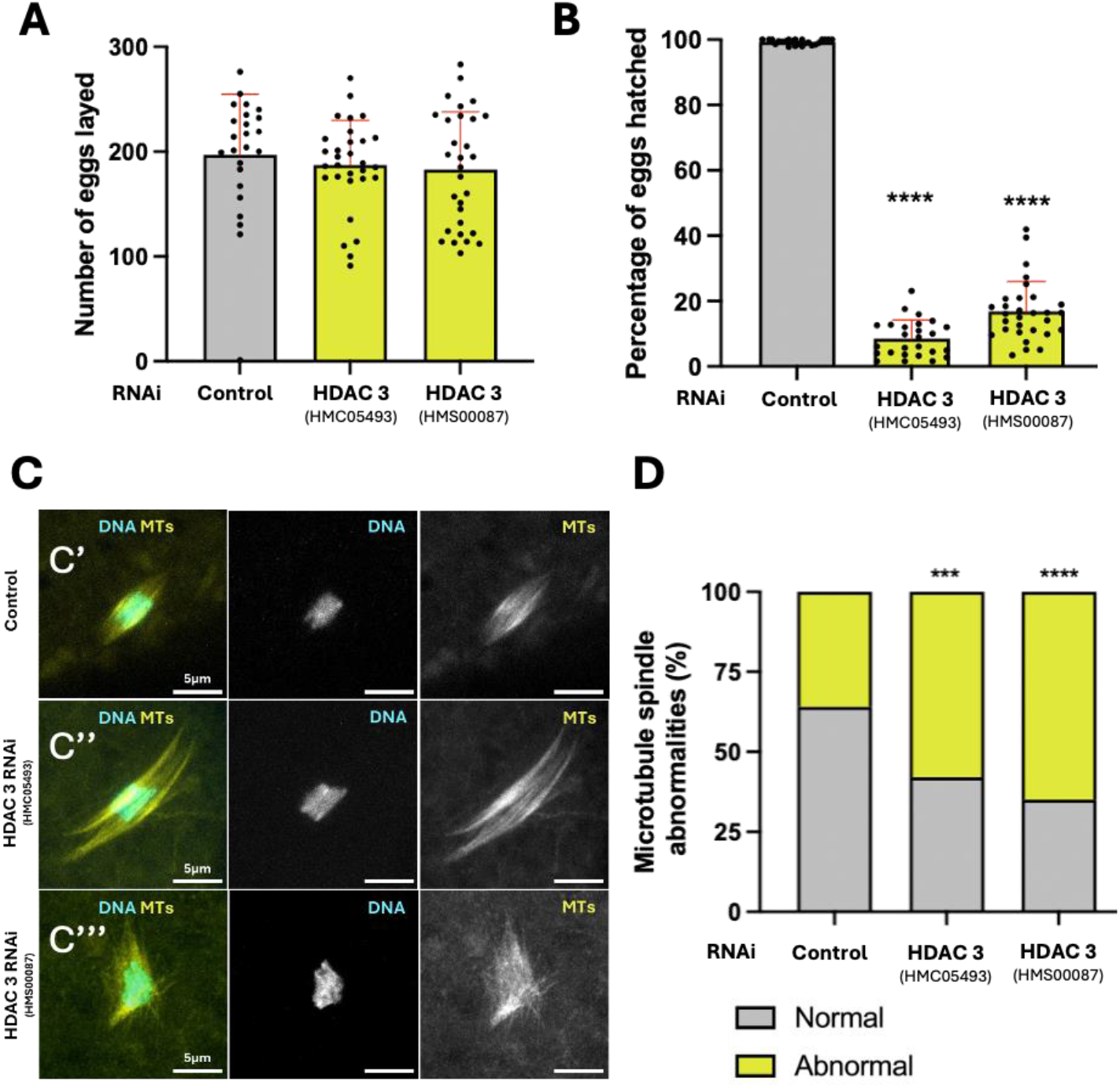
HDAC3 depletion leads to meiotic spindle defects and reduces female fertility. **(A-B)** Fertility assays perfomed in control (*UAS-mCherry RNAi)* and HDAC3 RNAi females. **(A)** Total number of eggs laid per female during 24h for control and HDAC3 RNAi. **(B)** Percentage of hatched eggs in the same conditions. Each dot represents one biological replicate (one cage); mean ± SEM are shown. Statistical analysis was performed using one-way ANOVA followed by Kruskal-Wallis’ multiple comparisons test comparing each RNAi condition to control. **** p < 0.0001. N = 3; n = 30. **(C-D)** Analysis of meiotic microtubule (MT) spindles in stage-14 oocytes from control (UAS-*mCherry* RNAi) and HDAC3 RNAi females. **(C)** Representative immunofluorescence images of meiotic spindles in stage-14 oocytes. Microtubules are labeled by fluorescently tagged Jupiter-GFP (yellow), and DNA is shown in cyan. Representative examples of morphologically normal (C′) and abnormal (C″, C‴) meiotic spindles are shown. Scale bar: 10 µm. **(D)** Quantification of meiotic spindle abnormalities (%). Statistical analysis was performed using one-way ANOVA followed by Kruskal–Wallis multiple-comparisons testing, with each RNAi condition compared to the control. ****p < 0.0001, ***p < 0.001. *N* = 3 independent biological replicates, with a minimum of 10 oocytes analyzed per experiment.

We further performed fertility assays in females in which HDAC3 was depleted from the germline. HDAC3 knockdown did not alter the number of eggs laid, indicating that earlier stages of oogenesis, including oocyte determination and mid-stage vitellogenesis, are largely unaffected (Fig. 4A). Consistent with previous work^41^, both RNAi lines showed a significant reduction in egg hatching rates, with the HDAC3^*HMC05493*^ line displaying the strongest effect (Fig. 4B), which parallels the severity of its karyosome phenotype (Fig. 3B,C). The reduced hatch rate is consistent with the spindle abnormalities observed during meiosis, as meiotic spindle defects can generate aneuploid oocytes with poor developmental potential^42,43^.

In oocytes, meiotic spindle assembly relies on chromatin-driven microtubule nucleation through the Ran-GTP pathway^44^. Both in *Drosophila* and mammals defects in chromatin condensation or higher-order chromosome structure directly cause abnormal spindle morphology in oocytes^42,45^, demonstrating that proper chromatin organization is required for assembly of a normal meiotic spindle. Consistent with this, in *Drosophila*, mutations or RNAi-induced depletion of proteins which disrupt the karyosome or chromosome condensation led to meiotic defects similar to the ones observed in HDAC3 depleted oocytes^7,36,37,46^.

Previous work in mice further supports a role of HDAC3 in meiotic progression as HDAC3-depleted mouse oocytes also displayed abnormal spindle morphology with disrupted chromosome alignment, and segregation errors. HDAC3 was reported to localize on meiotic spindles, and HDAC3-depleted oocytes showed increased microtubule acetylation, suggesting that HDAC3 may contribute to microtubule deacetylation^47^. This raises the possibility that the defects observed in the meiotic spindles upon HDAC3 depletion may be a consequence of microtubule hyperacetylation, however, expression of a non-acetylable tubulin only partially rescued the HDAC3 meiotic defects^48^, suggesting additional functions of HDAC3 in meiotic progression, independently of a function in MT deacetylation. Therefore, consistent with HDACs having multiple cellular targets, the meiotic defects observed in HDAC3-depleted oocytes likely arise from a combination of altered in chromatin architecture and additional abnormalities, including microtubule hyperacetylation. Taken together with previous studies in mouse oocytes, our findings suggest that HDAC3 performs a conserved role in ensuring proper meiotic progression across metazoans.

Our work highlights an important role for chromatin lactylation in oogenesis and oocyte developmental competence; however, we cannot exclude the possibility that the phenotypes observed upon depletion of HDAC3 or Nej/p300 arise partly from functions that extend beyond their role in regulating lactylation. Nevertheless, work in mammalian systems provides additional support for a functional connection between histone lactylation and oocyte developmental competence. Mouse oocytes also display a developmental dynamic regulation of Kla levels, with high levels of nuclear Kla also present at stages associated with highly compacted chromatin (germinal vesicle)^14,29^. Moreover, consistent with our observations, experimentally increasing histone lactylation levels in mouse oocytes caused changes in chromatin architecture and defects in metaphase-II spindle morphology and chromosome alignment^18^. Conversely, lowering lactate availability reduced Kla and impaired preimplantation development^14^. Together, these data suggest that proper control of histone lactylation during female gametogenesis is conserved across species and likely contributes to the acquisition of developmental competence.

Although depletion of HDAC3 or Nej/p300 each reduced histone lactylation levels, defects in karyosome morphology were only observed in the absence of HDAC3, and not Nej. One possibility is that these enzymes regulate distinct subsets of lactylated lysines, and that the sites controlled by HDAC3 are specifically required to maintain the compact chromatin architecture of the karyosome. Because we relied on a pan-lactylation antibody, our current analysis cannot distinguish residue-specific changes. Future work using site-specific Kla antibodies and mass spectrometry will be required to determine whether HDAC3 controls a functional subset of lactylation sites that are critical for chromatin architecture.

In *Drosophila* embryos, HDAC3 is known to contribute to transcriptional repression through non-catalytic mechanisms^49^. Therefore, it is also possible that the defects observed in HDAC3 depleted oocytes result from a combination of reduced histone lactylation together with impaired transcriptional repression.

We cannot exclude the possibility that the defects caused by HDAC3 depletion are associated with the disruption of its deacetylase activity, which could contribute to transcription deregulation. However, a previous screen for chromatin modifying enzymes did not detect any changes in the histone methylation marks associated with transcriptional activation or repression when HDAC3 was depleted using one of the RNAi lines used in our study^7^. This suggests that HDAC3 depletion does not significantly alter the transcription state under these conditions.

Together, our findings identify conserved chromatin-modifying enzymes that regulate histone lactylation in the female germ line and establish a foundation for future work aimed at dissecting the mechanistic and physiological roles of histone lactylation in oogenesis, meiotic fidelity, and fertility. Given that lactate metabolism is sensitive to nutrient and physiological state, our work raises the possibility that metabolic cues may interface with chromatin architecture and reproductive output through histone lactylation.

## MATERIALS AND METHODS

### Fly stocks and genetics

All *Drosophila* strains were maintained on standard medium at 25 °C using standard culture conditions. The following stocks were used: *w*^1118^ (control), Jupiter-GFP (Bloomington Drosophila Stock Center [BDSC] #6836), the maternal germline-specific V32-Gal4 driver (BDSC #7062), and UAS-Dcr2; nanos-Gal4 (BDSC #25751).

RNA interference (RNAi) lines used in this study included TRiP.HMC05493 (BDSC #64476) and TRiP.HMS00087 (BDSC #34778), targeting *hdac3*, and TRiP.HMS01570 (BDSC #36682), targeting *nej*. To achieve germline-specific depletion, RNAi lines were crossed to the UAS-Dcr2; nanos-Gal4 driver line, which is expressed in the female germ-line. An RNAi line targeting *mCherry* (BDSC #35785) was used as a negative control in RNAi experiments. For experiments analyzing meiotic spindles, RNAi lines were crossed to a single stock combining the V32-Gal4 driver with a Jupiter-GFP transgene, which labels microtubules, generated for this study.

### Ovaries and embryos immunostaining

Flies were fed with yeast for two days before dissection. Adult ovaries (10 ovary pairs per sample, per experiment) were dissected in BRB-80 buffer (80mM Pipes pH 6.8, 1mM MgCl2, 1 mM EGTA) with 1x protease inhibitors (Roche). After dissection, the ovaries were incubated in the same solution containing 1% Triton X-100 for 1h at room temperature (RT), followed by fixation in pre-cooled methanol at -20ºC for 15min. Three 5 min washes were performed in PBT (1× PBS, 0.1% Tween-20), followed by overnight permeabilization at 4 °C. Blocking was carried out in PBT with 2% BSA (Gibco). Primary antibodies were incubated overnight at 4ºC in PBS with 1% BSA under gentle agitation. After three 15 min washes in PBS with 1% BSA, secondary antibodies were incubated for 2h at RT in the same solution. DAPI (1:200; Roche) was added during the last 30 min to stain DNA. Samples were washed three times in PBS and mounted in Dako mounting medium (DAKO).

For the meiosis experiment, flies were on yeast for three days, and a modified fixation protocol was used. Adult ovaries were dissected in BRB-80 and permeabilized with 1% Triton X-100 in the same buffer for 1h at RT. Fixation was performed in PBS containing 4% PFA for 15 min at RT, followed by two 15 min washes in 0.1% PBST (PBS + 0.1 % Triton X-100), one additional wash in 0.1% PBST and a final wash in PBS. The ovaries were mounted using Vectashield with DAPI (Vector Laboratories).

For immunostaining analysis of *Drosophila* embryos, those were collected and fixed at 2-3 h of age. Embryos were dechorionated in 50% bleach for 5 min and fixed for 1 h with gentle shaking in 4 ml heptane, 0.287 ml 16% formaldehyde and 0,712 ml 1xPBS. Devitellization of the embryos was then performed by adding 8-9 ml of metanol (MeOH) and shaking vigorously for 1 min. Following rehydration in consecutive MeOH:PBS with 0,1% Tween-20 (PBT) solutions, increasing in ratio of PBT to MeOH, embryos were blocked in PBS containing 0,1% Tween-20, 1% BSA and 1% Donkey Serum (BBT + 1% Donkey Serum) at 4ºC O/N. Primary antibody incubations were carried in BBT + 1% Donkey Serum at 4ºC O/N. Embryos were then thoroughly washed in PBT and reblocked in BBT + 1% Donkey Serum for 1 h at room temperature, before the incubation of the secondary antibody for 2 h at room temperature. Secondary antibody incubation was also carried out in BBT + 1% Donkey Serum. DNA was stained with DAPI at 1:200 for another 30 min following secondary antibody incubation. Embryos were extensively washed in PBT and mounted in Fluorescent Mounting Medium.

### Antibodies

Primary antibodies: rabbit anti-L-Lactyllysine (1:500; PTM-Biolabs; PTM-1401) and mouse anti-Orb (clones 4H8 and 6H4, 1:30 dilution each, Developmental Studies Hybridoma Bank). Secondary antibodies: Alexa Fluor 488 anti-mouse (1:500; ThermoFisher, A21202) and Alexa Fluor 647 anti-rabbit (1:500; ThermoFisher, A31573).

### Imaging, analysis and quantification

Egg chambers were imaged on a Zeiss LSM980 confocal microscope using a 63×/1.4 NA oil-immersion objective and ZEN Blue 3.3 software. Serial optical sections were acquired every 0.30 µm. Imaging parameters were held constant across all samples and biological replicates within each experiment to allow quantitative comparisons.

Background signal was estimated by measuring three regions of interest per image; average background intensity was subtracted before further analysis. Images were converted to sum-intensity projections. An outline was drawn around each oocyte chromatin region, and nuclear lactylation intensity was measured in arbitrary units (a.u.) using ImageJ (NIH). Values were normalized to the mean intensity of the control group. At least three independent biological replicates were analyzed for each condition.

For DNA condensation quantification, a qualitative classification was first performed, categorizing oocyte chromatin as condensed, starting to decondense, or decondensed. Z-stacks were then projected into maximum-intensity images, and nuclear morphology was quantified in ImageJ by measuring area, perimeter, circularity, and solidity. Circularity was calculated as 4π × (area/perimeter^2^), ranging from 0 (elongated) to 1 (perfect circle). Solidity was calculated as (area/convex area), ranging from 0 (irregular) to 1 (compact). A minimum of three independent experiments was performed for each condition.

### Fertility assays

For fertility assays, eight well-fed females (7 days old) of each RNAi genotype were crossed with four *w*^1118^ males in cages containing agar plates supplemented with apple juice. For each genotype, ten independent cages were analyzed per experiment. Eggs laid within a 24 h period were counted, after which plates were maintained at 25 °C for an additional 2 da ys and the number of hatched larvae was scored. The percentage of egg hatching was calculated as the ratio of hatched larvae to total eggs laid. The entire experiment was repeated three times using independent biological replicates, with newly collected females and males for each replicate.

### Statistics

At least three independent biological replicates were performed for all experiments. The number of egg chambers and oocytes analyzed in each experiment is indicated in the figure legends. The statistical tests used for each analysis are indicated in the corresponding figure legends. Statistical analyses were performed using GraphPad Prism 10.

## ACKNOWLEDGMENTS

We thank all members of the Cytoskeleton in Development and Disease lab. (NOVA Medical School, NMS) and the Cellular Plasticity in Regeneration and Cancer Lab. (NOVA Institute for Medical Systems Biology), both at NOVA University Lisbon, for their feedback and insightful discussions that contributed to the development of this work. We are grateful to Rui Martinho and Catarina Homem for valuable scientific discussions. We thank Mónica Bettencourt-Dias (Gulbenkian Institute for Molecular Medicine, Oeiras, Portugal) for sharing fly stocks.

We acknowledge the support of the Fly Facility and the Advanced Imaging Facility at NMS for infrastructure and technical support.

I.G. is supported by a doctoral fellowship from the Portuguese Foundation for Science and Technology (FCT; UI/BD/154334/2022). A.P.M. is supported by an individual CEECIND contract from FCT (2020.02842.CEECIND).

This work was supported by the Research Unit iNOVA4Health (UID/4462/2025) and by the Associated Laboratory LS4FUTURE (LA/P/0087/2020), all financially supported by Fundação para a Ciência e Tecnologia/Ministério da Educação, Ciência e Inovação.

## REFERENCES

1. Conti, M. & Franciosi, F. Acquisition of oocyte competence to develop as an embryo: integrated nuclear and cytoplasmic events. Hum Reprod Update 24, 245–266 (2018).

2. Cabrita, B. & Martinho, R. G. Genetic and Epigenetic Regulation of Drosophila Oocyte Determination. Journal of Developmental Biology 2023, Vol. 11, Page 21 11, 21 (2023).

3. Bannister, A. J. & Kouzarides, T. Regulation of chromatin by histone modifications. Cell Research 2011 21:3 21, 381–395 (2011).

4. Allis, C. D. & Jenuwein, T. The molecular hallmarks of epigenetic control. Nat Rev Genet 17, 487–500 (2016).

5. He, M., Zhang, T., Yang, Y. & Wang, C. Mechanisms of Oocyte Maturation and Related Epigenetic Regulation. Front Cell Dev Biol 9, (2021).

6. De La Fuente, R. Chromatin modifications in the germinal vesicle (GV) of mammalian oocytes. Dev Biol 292, 1–12 (2006).

7. Navarro-Costa, P. et al. Early programming of the oocyte epigenome temporally controls late prophase I transcription and chromatin remodelling. Nature Communications 2016 7:1 7, 1–15 (2016).

8. Sindik, N., Pereza, N., Sanja, · & Pavlić, D. Epigenetics of oogenesis. 311, 183–190 (2025).

9. He, M., Zhang, T., Yang, Y. & Wang, C. Mechanisms of Oocyte Maturation and Related Epigenetic Regulation. Front Cell Dev Biol 9, 654028 (2021).

10. Zhang, D. et al. Metabolic regulation of gene expression by histone lactylation. Nature 2019 574:7779 574, 575–580 (2019).

11. Galle, E. et al. H3K18 lactylation marks tissue-specific active enhancers. Genome Biol 23, (2022).

12. Meng, X., Baine, J. M., Yan, T. & Wang, S. Comprehensive Analysis of Lysine Lactylation in Rice (Oryza sativa) Grains. J Agric Food Chem 69, 8287–8297 (2021).

13. Merkuri, F., Rothstein, M. & Simoes-Costa, M. Histone lactylation couples cellular metabolism with developmental gene regulatory networks. Nat Commun 15, (2024).

14. Yang, W. et al. Hypoxic in vitro culture reduces histone lactylation and impairs pre-implantation embryonic development in mice. Epigenetics Chromatin 14, 1–15 (2021).

15. Li, H., Sun, L., Gao, P. & Hu, H. Lactylation in cancer: Current understanding and challenges. Cancer Cell 42, 1803–1807 (2024).

16. Pan, R. Y. et al. Positive feedback regulation of microglial glucose metabolism by histone H4 lysine 12 lactylation in Alzheimer’s disease. Cell Metab 34, 634–648.e6 (2022).

17. Hagihara, H. et al. Protein lactylation induced by neural excitation. Cell Rep 37, (2021).

18. Lin, J. et al. Overexpression of Tfap2a in Mouse Oocytes Impaired Spindle and Chromosome Organization. International Journal of Molecular Sciences 2022, Vol. 23, Page 14376 23, 14376 (2022).

19. Zhang, X., Liu, Y. & Wang, N. Dynamic changes in histone lysine lactylation during meiosis prophase I in mouse spermatogenesis. Proc Natl Acad Sci U S A 122, (2025).

20. Hayashi, Y. et al. Comprehensive observation of histone lysine lactylation during gametogenesis of Drosophila melanogaster. Developmental Dynamics 10.1002/DVDY.70010 (2025) doi:10.1002/DVDY.70010.

21. Mahowald, A. P. & Tiefert, M. Fine structural changes in the Drosophila oocyte nucleus during a short period of RNA synthesis - An autoradiographic and ultrastructural study of RNA synthesis in the oocyte nucleus of Drosophila. Wilhelm Roux Arch Entwickl Mech Org 165, 8–25 (1970).

22. Bogolyubov, D. S. Karyosphere (Karyosome): A Peculiar Structure of the Oocyte Nucleus. Int Rev Cell Mol Biol 337, 1–48 (2018).

23. King, R. C. The Meiotic Behavior of the Drosophila Oocyte. Int Rev Cytol 28, 125–168 (1970).

24. Lantz, V., Chang, J. S., Horabin, J. I., Bopp, D. & Schedl, P. The Drosophila orb RNA-binding protein is required for the formation of the egg chamber and establishment of polarity. Genes Dev 8, 598–613 (1994).

25. Peng, X. & Du, J. Histone and non-histone lactylation: molecular mechanisms, biological functions, diseases, and therapeutic targets. Molecular biomedicine 6, (2025).

26. Wang, J. et al. Mechanism and application of lactylation in cancers. Cell & Bioscience 2025 15:1 15, 76– (2025).

27. Dolsten, G. A. et al. 3D chromatin structures precede genome activation in Drosophila embryogenesis. Cell genomics 5, (2025).

28. Ciabrelli, F., Atinbayeva, N., Pane, A. & Iovino, N. Epigenetic inheritance and gene expression regulation in early Drosophila embryos. EMBO Reports 2024 25:10 25, 4131–4152 (2024).

29. Yang, D. et al. Histone Lactylation Is Involved in Mouse Oocyte Maturation and Embryo Development. Int J Mol Sci 25, (2024).

30. Vidaurre, V. & Chen, X. Epigenetic Regulation of Drosophila Germline Stem Cell Maintenance and Differentiation. Dev Biol 473, 105 (2021).

31. Hu, Y. et al. Lactylation: the novel histone modification influence on gene expression, protein function, and disease. Clin Epigenetics 16, 72– (2024).

32. Gonzatti, M. B. et al. Class I histone deacetylases catalyze lysine lactylation. Journal of Biological Chemistry 301, 110602 (2025).

33. Yang, K. et al. Lactate promotes macrophage HMGB1 lactylation, acetylation, and exosomal release in polymicrobial sepsis. Cell Death & Differentiation 2021 29:1 29, 133–146 (2021).

34. Fan, M. et al. Lactate promotes endothelial-to-mesenchymal transition via Snail1 lactylation after myocardial infarction. Sci Adv 9, (2023).

35. Moreno-Yruela, C. et al. Class I histone deacetylases (HDAC1-3) are histone lysine delactylases. Sci Adv 8, 6696 (2022).

36. Sun, S. et al. Metabolic regulation of cytoskeleton functions by HDAC6-catalyzed α-tubulin lactylation. Nature Communications 2024 15:1 15, 8377– (2024).

37. Zhaunova, L., Ohkura, H. & Breuer, M. Kdm5/Lid Regulates Chromosome Architecture in Meiotic Prophase I Independently of Its Histone Demethylase Activity. PLoS Genet 12, e1006241 (2016).

38. Ivanovska, I., Khandan, T., Ito, T. & Orr-Weaver, T. L. A histone code in meiosis: the histone kinase, NHK-1, is required for proper chromosomal architecture in Drosophila oocytes. Genes Dev 19, 2571–2582 (2005).

39. Lancaster, O. M., Cullen, C. F. & Ohkura, H. NHK-1 phosphorylates BAF to allow karyosome formation in the Drosophila oocyte nucleus. J Cell Biol 179, 817–824 (2007).

40. Lowe, N. et al. Analysis of the expression patterns, Subcellular localisations and interaction partners of drosophila proteins using a pigp protein trap library. Development (Cambridge) 141, 3994–4005 (2014).

41. Karpova, N., Bobinnec, Y., Fouix, S., Huitorel, P. & Debec, A. Jupiter, a newDrosophila protein associated with microtubules. Cell Motil Cytoskeleton 63, 301–312 (2006).

42. Tang, M. et al. Separation of transcriptional repressor and activator functions in Drosophila HDAC3. Development (Cambridge) 150, (2023).

43. Radford, S. J., Nguyen, A. L., Schindler, K. & McKim, K. S. The chromosomal basis of meiotic acentrosomal spindle assembly and function in oocytes. Chromosoma 2016 126:3 126, 351–364 (2016).

44. Hughes, S. E., Miller, D. E., Miller, A. L. & Hawley, R. S. Female Meiosis: Synapsis, Recombination, and Segregation in Drosophila melanogaster. Genetics 208, 875 (2018).

45. Cesario, J. & McKim, K. S. RanGTP is required for meiotic spindle organization and the initiation of embryonic development in Drosophila. J Cell Sci 124, 3797–3810 (2011).

46. Resnick, T. D. et al. Mutations in the chromosomal passenger complex and the condensin complex differentially affect synaptonemal complex disassembly and metaphase I configuration in Drosophila female meiosis. Genetics 181, 875–887 (2009).

47. Loh, B. J., Cullen, C. F., Vogt, N. & Ohkura, H. The conserved kinase SRPK regulates karyosome formation and spindle microtubule assembly in Drosophila oocytes. J Cell Sci 125, 4457–4462 (2012).

48. Li, X. et al. HDAC3 promotes meiotic apparatus assembly in mouse oocytes by modulating tubulin acetylation. Development 144, 3789–3797 (2017).

49. He, Y., Li, X., Gao, M., Liu, H. & Gu, L. Loss of HDAC3 contributes to meiotic defects in aged oocytes. Aging Cell 18, e13036 (2019).

50. Lewandowski, S. L., Janardhan, H. P. & Trivedi, C. M. Histone Deacetylase 3 Coordinates Deacetylase-independent Epigenetic Silencing of Transforming Growth Factor-β1 (TGF-β1) to Orchestrate Second Heart Field Development. J Biol Chem 290, 27067–27089 (2015).

