## Supplementary material for "A role for HDAC3 in regulating histone lactylation and maintaining oocyte chromatin architecture and fertility": Gomes et al., 2026_SUPP_DATA_final.docx

**Supplementary Figure 1**

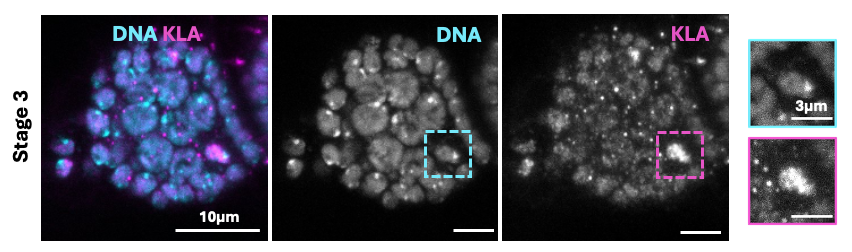

**Supplementary Figure 1: Histone lactylation increases abruptly at stage 3 in *Drosophila* oocytes.** Representative immunofluorescence images of stage 3 Drosophila oocytes showing histone lysine lactylation. Kla in magenta and DNA in cyan. Merged and individual channels are displayed. Insets show higher-magnification views of oocyte nuclei. Scale bars: 10 µm and 3 µm (insets).

**Supplementary Figure 2:**

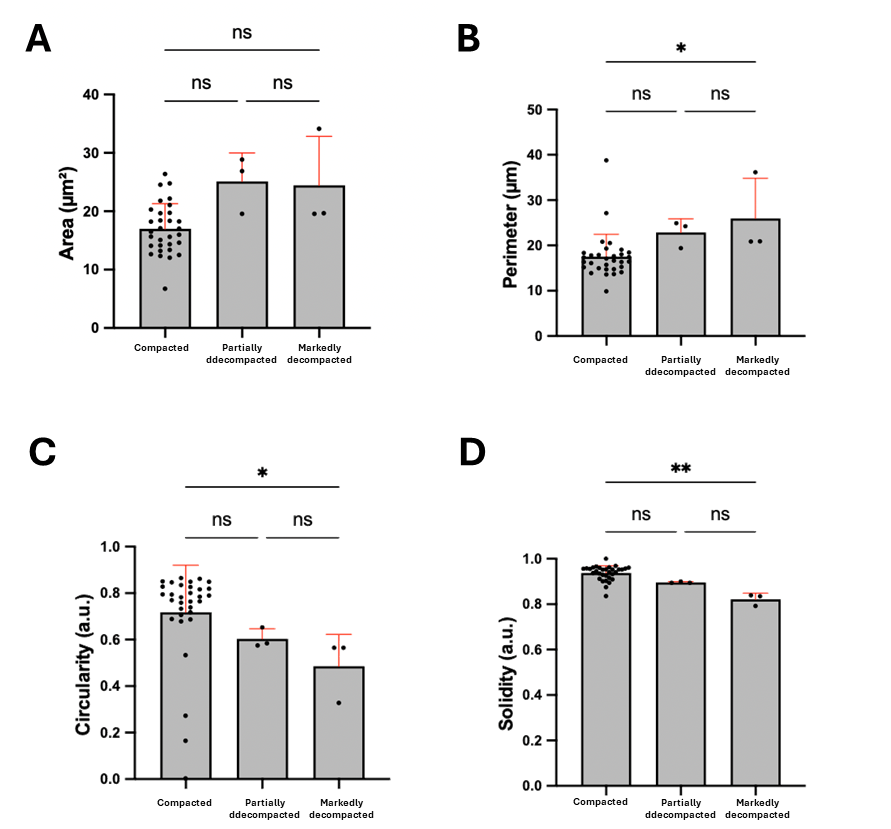

**Supplementary Figure 2:** **Quantification of karyosome geometric parameters in control oocytes.** **(A–D)** Quantification of nuclear area (µm²), perimeter (µm), circularity (4π × area / perimeter², range 0–1, with 1 = perfect circle), and solidity (area / convex area, range 0–1, with 1 = compact nucleus) in control oocytes (*mCherry* RNAi). Oocytes were first classified into three karyosome morphology categories as defined in Fig. 3, and geometric parameters were then measured independently for each category to assess differences between groups. Each dot represents one oocyte nucleus; mean ± SEM are shown. Statistical analysis was performed using one-way ANOVA followed by Kruskal-Wallis’ multiple comparisons test comparing each RNAi condition to control. ** p < 0.01, * p < 0.05, ns = not significant. N = 3; minimum of 10 oocytes per experiment.

**Supplementary Figure 3:**

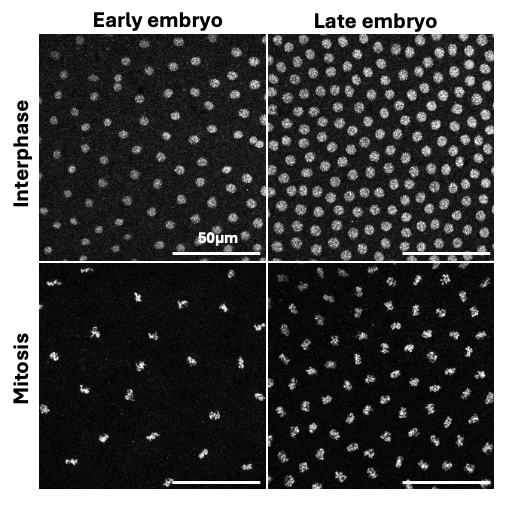

**Supplementary Figure 3:** **Histone lactylation is present in the embryo, at early and later mitotic cycles, in both mitotic and interphasic nuclei.** Representative images of control Drosophila embryos immunostained for lysin histone lactylation (Kla). Scale bar: 50µm.
